# Efficient Gene Targeting in Maize using Inducible CRISPR-Cas9 and Marker-Free Donor Template

**DOI:** 10.1101/2020.05.13.093575

**Authors:** Pierluigi Barone, Emily Wu, Brian Lenderts, Ajith Anand, William Gordon-Kamm, Sergei Svitashev, Sandeep Kumar

**Author notes:** These authors contributed equally to this work (Pierluigi Barone, Emily Wu).

## Abstract

CRISPR-Cas9 is a powerful tool for generating targeted mutations and genomic deletions. However, precise gene insertion or sequence replacement remains a major hurdle before application of CRISPR-Cas9 technology is fully realized in plant breeding. Here we report high frequency, selectable marker-free intra-genomic gene targeting (GT) in maize. Heat shock-inducible Cas9 was used for generating targeted double-strand breaks (DSBs) and simultaneous mobilization of the donor template from pre-integrated T-DNA. The construct was designed such that release of the donor template and subsequent DNA repair activated expression of the selectable marker gene within the donor locus. This approach generated up to 4.7% targeted insertion of the donor sequence into the target locus in T0 plants, with up to 86% detected donor template release and 99% mutation rate were observed at the donor loci and the genomic target site, respectively. Unlike previous *in planta* or intra-genomic homologous recombination reports, that required multiple generations and extensive screening, our method provides non-chimeric, heritable GT in the T0 generation.

## Introduction

CRISPR (clustered regularly interspaced short palindromic repeats)-Cas9^1^ has emerged as a robust and versatile genome editing tool to generate targeted DNA double-strand breaks^1–5^. This approach overcomes the limitations of conventional breeding and accelerates development of improved plant varieties. Inherent cellular DSB repair mechanisms rely on either non-homologous end joining (NHEJ) or homologous recombination (HR)^6^. NHEJ is an error-prone repair pathway that involves ligation of non-homologous sequences or sequences with micro-homologies, often resulting in small insertions or deletions (indels) at the DSB site. HR requires a DNA repair template with sequences homologous to those flanking the genomic DSB site, leading to precise repair of the DSB. NHEJ is the primary DNA repair pathway in somatic cells, while HR is observed with much lower frequencies and predominantly occurs during S and G2 phases of the cell cycle^7^.

CRISPR-Cas9-mediated genome editing can be used to improve agronomic characteristics, introduce targeted disease resistance^8–12^, and speed-up domestication for development of new beneficial plant varieties^13^. Some traits that depend on endogenous gene mutation can be generated through NHEJ-mediated DNA repair ^14–18^. Other traits require precise genome editing and need gene targeting (GT) via HR to make large insertions or allele replacements^19,20^. While high frequency NHEJ-mediated gene mutations (30–100%) have been obtained in plants^21^, precise GT via HR remains exceedingly low and a major improvement is needed for routine application. Targeted mutagenesis relies on NHEJ-mediated repair of DSBs without any repair template, and therefore has become routine in plants that are amenable to transformation and regeneration. HR-mediated GT requires simultaneous creation of targeted DSB(s) and supply of a donor DNA repair template containing flanking regions of homology (also referred to as homology arms)^22^. While targeted DSBs can render a desired genomic site accessible to a donor repair template, the DSB does not prevent ectopic integration of a donor sequence elsewhere in the genome. Therefore, high frequency random integration of donor template exacerbated by limitations in transformation and regeneration processes makes HR-mediated GT in plants extremely inefficient. Positive selection for GT using a promoter trap or complementation of a non-functional selectable marker in the target locus has been used successfully to avoid regeneration of plants with random integration of the donor template^23–26^. In addition, *in planta*^27–29^ and intra-genomic homologous recombination (IGHR)^22,30^ approaches have been used to address plant transformation inefficiency. Since IGHR describes GT in both tissue culture and whole plant (*in planta*), the term IGHR in this manuscript conforms to this broader interpretation. The IGHR approach utilizes stable integration of donor sequence, followed by extensive screening of progeny plants to detect rare GT in subsequent generations. Long generation time and low efficiency of GT have been two major bottlenecks for routine application of this method in plants.

Here we report a method for high frequency GT in maize, where the resultant HR-modified target locus is potentially selectable marker-free. Using heat shock inducible Cas9 expression, donor template was mobilized intra-genomically during early stages of maize transformation. Consequently, Cas9-mediated release of the donor template activated a selectable marker gene *Hra*^31^ (**H**ighly herbicide **R**esistant ***A**ls*), which allowed preferential regeneration of events from cells where the donor template has been excised. Even though the donor template contained *NptII* (*neomycin phosphotransferase II*) gene, G418 (Geneticin) selection was not used during plant regeneration. Instead, the *NptII* sequence represented the gene of interest being introduced into the target locus. This approach generated up to 4.7% putative GT positive T0 plants, where 86% of these plants had donor template release and 99% mutation rate at the target site. Analyses of T1 progeny confirmed Mendelian inheritance of GT events. For the majority of the GT-positive events, randomly integrated T-DNA segregated independently of the GT locus resulting in T-DNA-free GT-positive T1 progeny. Analysis of T1 plants also revealed two bi-allelic GT T0 progenitor plants. To our knowledge, this is the first report describing robust gene targeting of selectable marker-free donor template in a crop plant. Unlike previous *in planta*^27,29^ and intra-genomic methods, this new approach generates GT directly in the T0 plants without going through time-consuming crossing and labor-intensive screening over multiple generations.

## Results

### Rationale for a new intra-genomic Gene Targeting (GT) system

An overview of the High-frequency Intra-Genomic Homologous Recombination (High-IGHR) system used in this study is outlined in Figure 1. The T-DNA construct (Figure 1A) contained donor DNA repair template with homology arms, HR1 and HR2, which was flanked by two Cas9 cut sites matching the genomic target site. The donor DNA repair template was placed between the promoter and *Hra* coding sequence, rendering the selectable marker non-functional. The T-DNA construct also contained a single guide RNA and an inducible Cas9 expression cassette. Following T-DNA transfer, the construct was randomly integrated into the genome, creating the donor locus (Figure 1B). Upon subsequent induction of Cas9 expression, two DSBs were generated in the donor locus outside each of the homology arms leading to release of the donor DNA repair template (Figure 1C), referred to as “donor release”. In addition, a DSB was generated at the genomic target locus (Figure 1D), which could then be repaired via HR using the extra-chromosomal (free) donor template, resulting in GT (Figure 1E). Finally, donor release activated the selectable marker gene by bringing the *Hra* coding sequence into functional proximity of the upstream promoter in the repaired donor locus (Figure 1F). *Hra* activation provided preferential selection for plant regeneration from cells with released donor template and potential GT (Figure 1G).

**Figure 1.**
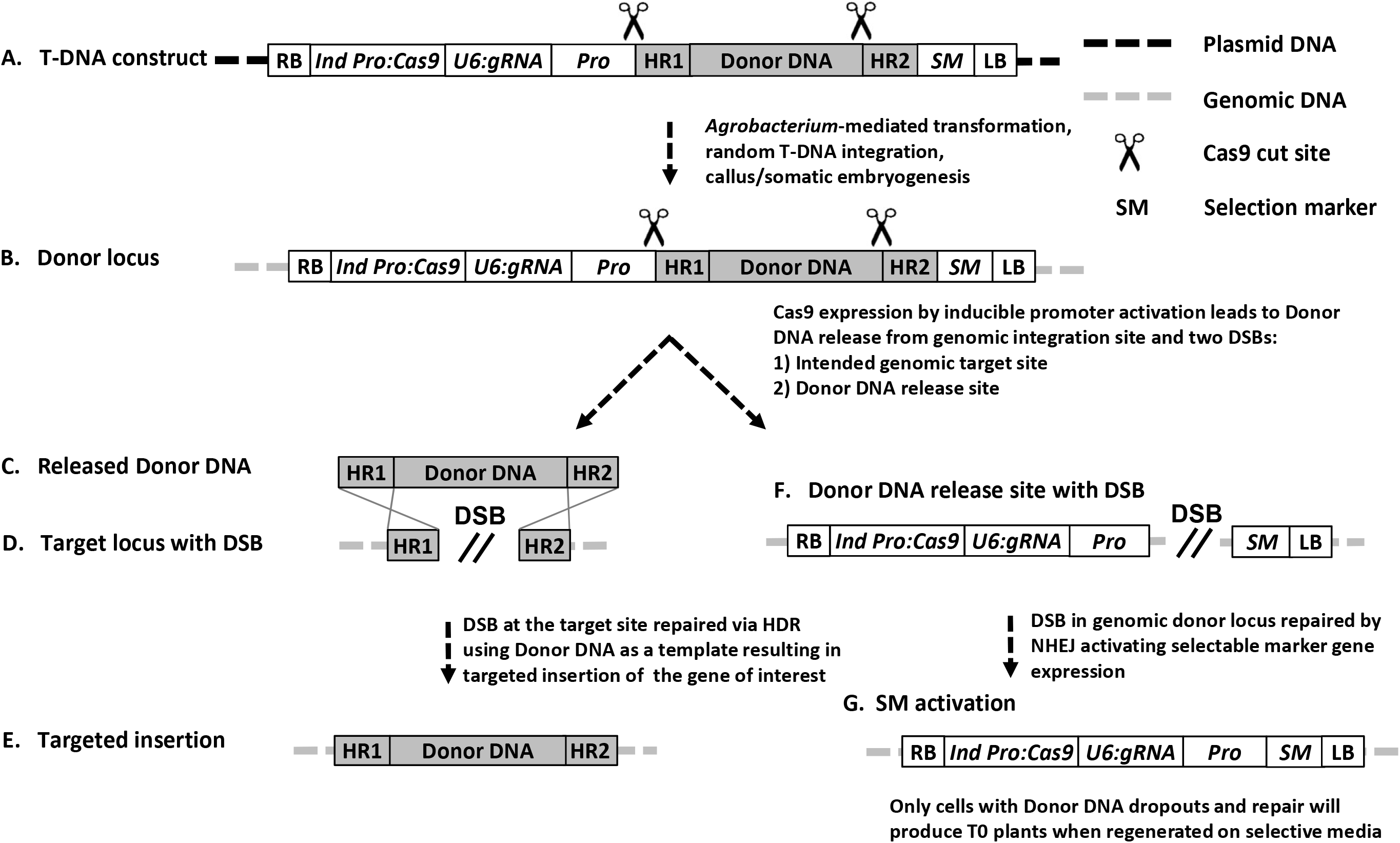
Inducible Cas9 and selectable marker activation system for HR-mediated intragenomic GT in maize. **A**. Schematic description of T-DNA construct containing Cas9 driven by an inducible promoter (*Ind Pro*), guide RNA expressed by an U6 promoter, donor flanked by homology arms (HR1 and HR2) and Cas9 cut sites (indicated by scissors) positioned between selectable marker (SM) and upstream promoter such that selectable marker is non-functional in this configuration. **B**. Stable random integration of T-DNA into the plant genome. **C & D**. Induction of Cas9 expression releases donor DNA repair template and simultaneously generates DSB at the target site, which is repaired using HR. **E**. HR-mediated repair at target site leads to targeted insertion of the gene of interest. **F & G**. The DSB at donor locus repaired by NHEJ resulting in activation of functional selectable marker.

### Intra-genomic GT using inducible Cas9

A plant transformation construct containing heat shock inducible *Hsp26*^32^ promoter-driven *Cas9* was used for intra-genomic GT (Figure 2A). Donor template in the construct included a *NptII* expression cassette flanked by homology arms specific to a maize genomic target site on chromosome 1 designated here as TS45^33^. The donor template was positioned between the coding sequence of an *Hra* gene (selectable marker) and the upstream *Sb-Als* promoter (Supplementary Table 1), preventing *Hra* expression in this starting configuration. The construct also contained two morphogenic genes, maize phospholipid transferase gene promoter (*Pltp*)-driven *Bbm* and maize auxin-inducible promoter (*Axig1*)-driven *Wus2*^34^.

**Figure 2.**
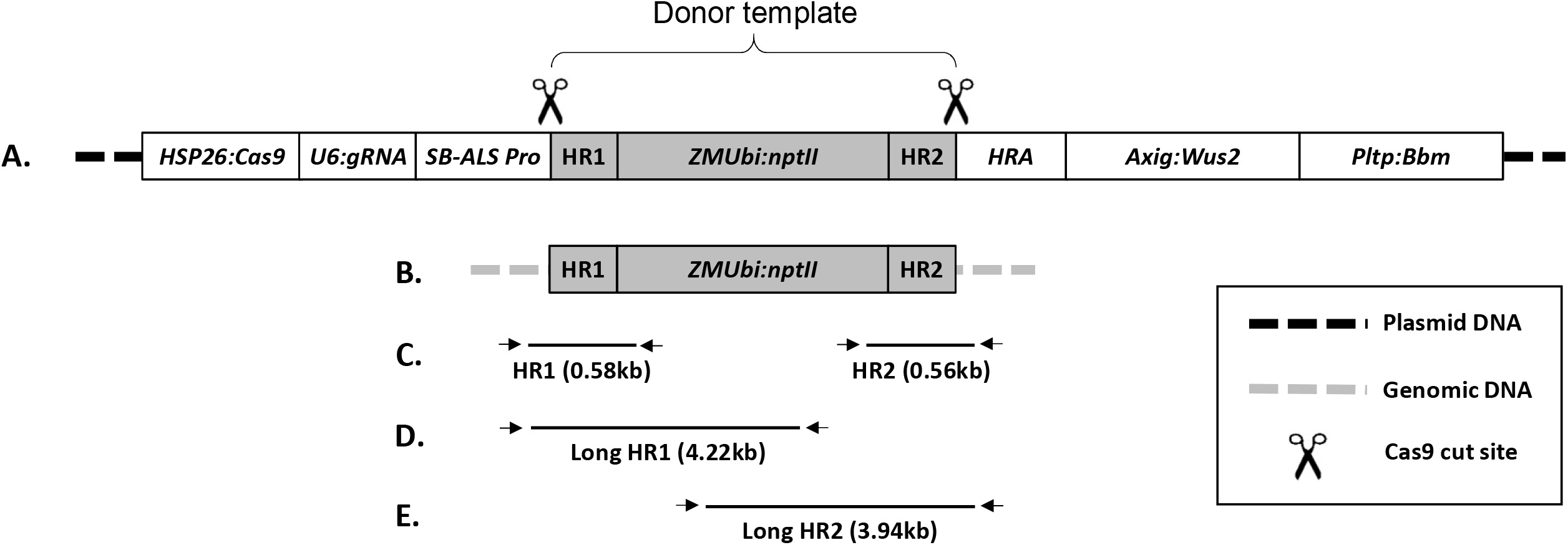
Schematic description of the T-DNA construct and PCR analysis design to detect gene targeting. **A**. Construct contains heat shock-inducible promoter 26 (HSP26) driven *Cas9*, maize U6 promoter to express guide RNA. Donor template containing *ZMUbi:nptII* gene expression cassette flanked by homology arms (411 bp HR1 and 422 bp HR2) and Cas9 cut sites (shown as scissors) positioned between sorghum ALS promoter and selectable marker gene HRA (herbicide resistant ALS) making. The construct also has morphogenic genes – *Axig* promoter driven *Wus2* and *Bbm* controlled by *Pltp* promoter. **B**. Putative perfect GT event. **C**. Diagnostic HR1 and HR2 junction PCR used for GT detection in T0 plants. **D & E**. Long overlapping PCR used for GT confirmation in T1 plants.

A total of 1213 maize immature embryos were transformed and subjected to heat shock treatments as depicted in Figure 3. One hundred and eleven embryos were used in the control treatment where no heat shock was performed. The remaining 1102 embryos were split into two experimental groups of 551 embryos each. These two groups were subjected to heat shock treatment at either 4 (HS4) or 11 (HS11) days after *Agrobacterium* infection (Figure 3). After initial heat shock treatment (HS4 or HS11), a second heat shock was applied the following day. Subsequently, the explants from the HS4 treatment were incubated on resting media for 7 days without selection. No resting period was given to treatment HS11 to ensure explants from both treatments and non-HS control were moved to HRA selection media 12 days after *Agrobacterium* infection. T0 plants regeneration was performed on HRA selection media as described in Materials and Methods. A total of 328 and 127 plants were regenerated from the HS4 and HS11 group, respectively (Table 1). Only four plants were regenerated from the control group of embryos not subjected to the heat shock, indicating tight regulation of Cas9 expression under *Hsp26* promoter.

**Figure 3.**
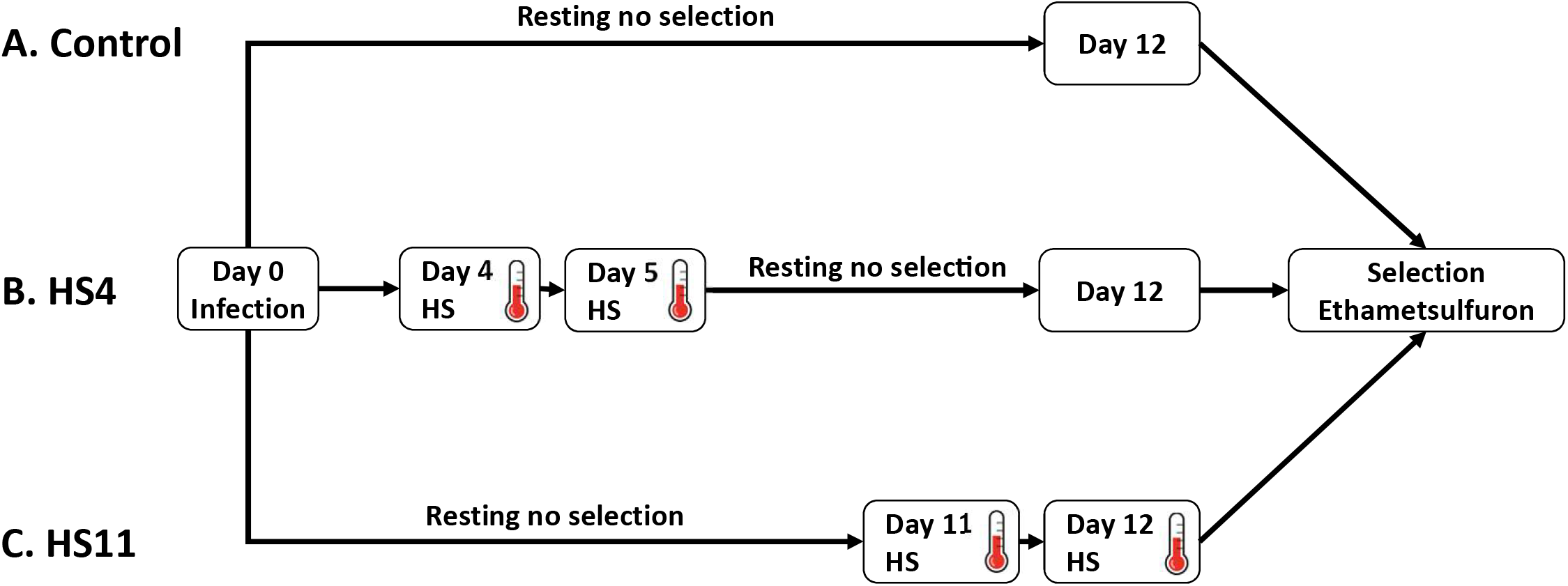
Work flow of heat shock treatments to induce Cas9 expression. **A**. Control treatment was not subjected to heat shock treatment. After *Agrobacterium* infection at day 0, explants were transferred to no selection medium (resting) until day 12. **B**. Two consecutive heat shock treatments were given at day 4 and day 5 for HS4 treatment. **C**. Two consecutive heat shock treatments were given at day 11 and day 12 for HS11 treatment. Explants from all three treatments were moved to selection medium with Ethametsulfuron 12 days after *Agrobacterium* infection.

**Table 1.**
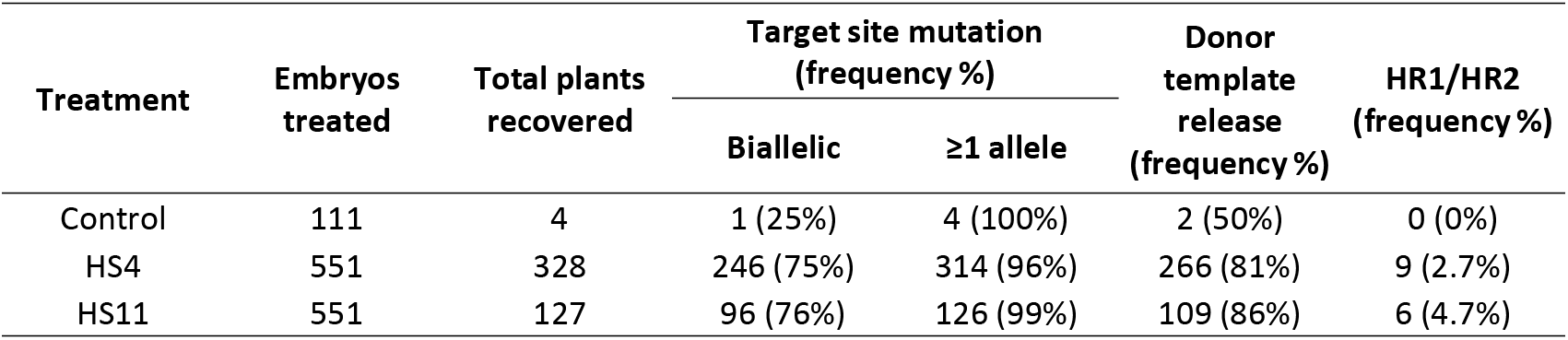
Molecular analyses of T0 plants for target site mutation and GT frequency

| Treatment | Embryos treated | Total plants recovered | Target site mutation (frequency %) |  | Donor template release (frequency %) | HR1/HR2 (frequency %) |
| --- | --- | --- | --- | --- | --- | --- |
|  |  |  | Biallelic | ≥1 allele |  |  |
| Control | 111 | 4 | 1 (25%) | 4 (100%) | 2 (50%) | 0 (0%) |
| HS4 | 551 | 328 | 246 (75%) | 314 (96%) | 266 (81%) | 9 (2.7%) |
| HS11 | 551 | 127 | 96 (76%) | 126 (99%) | 109 (86%) | 6 (4.7%) |

HR-mediated GT was detected by diagnostic HR1 and HR2 junction qPCR as depicted in Figure 2B-C for T0 plant and long overlapping PCR (Figure 2D-E) for T1 plant analysis. For plants recovered from the HS4 treatment, junction-PCR products for both HR1 and HR2 junctions were observed in 9 out of 328 T0 plants (2.7%), while for the HS11 treatment, 6 out of 127 plants (4.7%) were HR1/HR2 junction-PCR positive. Mutation analysis of the target site revealed up to 99% of T0 plants containing mutations in at least one allele, while >75% of the plants were observed with bi-allelic mutations (Table 1). High frequencies of donor template release were observed from plants regenerated using both HS4 and HS11 heat shock treatments (81% and 86%, respectively).

### Inheritance and segregation analysis of GT plants

To study inheritance and segregation of GT, T0 regenerants were crossed to wild-type plants to generate T1 progeny. Forty-seven T1 plants from each of the 13 GT-positive (representing both heat shock treatments) and three GT-negative (control group) T0 regenerants were analyzed for the presence of GT and T-DNA components. To confirm GT and validate integrity of the insertion, long overlapping 5’- and 3’-end PCR was performed as depicted in Figure 2D-E. Progeny from all 13 GT-positive T0 plants conformed to expected Mendelian inheritance of the modified targeted loci. Out of 47 T1 plants analyzed for each T0 regenerant, approximately 50% were GT-positive for progeny from 11 T0 plants, consistent with stable Mendelian inheritance of a mono-allelic GT locus (Table 2). From the remaining two T0 plants (plants 5 and 10), all progenies were GT-positive, which is consistent with bi-allelic GT at the T0 stage (Table 2). All T1 plants were also tested for the presence of randomly integrated T-DNA components including gRNA, *Hra*, and sequences proximal to T-DNA borders. HR1 and HR2 positive and T-DNA-free GT progenies were obtained from 12 GT-positive T0 plants. No T-DNA-free T1 plants were obtained only for one bi-allelic GT T0 plant (plant 10). As expected, no positive GT T1 plants were obtained from the three control T0 regenerants that tested negative for GT (plants 14, 15, and 16). Sequencing of PCR products from selected T1 progenies from each group confirmed expected targeted insertion in 7 out of 12 T0 plants. Deletion of approximately 600 bp of the ZMUBI intron of donor sequence was observed in progeny from two T0 plants (plant 4 and 11), while three T0 plants (5, 9, and 13) contained an approximately 400 bp duplication of the HR2 homology sequence at the 3’ end of the donor template (Table 2).

**Table 2.** Inheritance and segregation of GT in T1 generation

| T0 plant |  |  | T1 plants |  |  | Sequence confirmation |
| --- | --- | --- | --- | --- | --- | --- |
| ID Number | Treatment | Transgene copy number | CRISPR-Cas9 positive | HR1/2 positive | CRISPR Cas9-free & HR1/2 positive |  |
| 1 | HS4 | 2 | 40* | 21 | 2* | Yes |
| 2 |  | 2 | 17 | 21 | 13 | Yes |
| 3 |  | 2 | 24 | 25 | 14 | Yes |
| 4 |  | 2 | 18 | 20 | 13 | No<sup>a</sup> |
| 5 |  | 3 | 30 | 47* | 11 | No<sup>b</sup> |
| 6 |  | 1 | 17 | 24 | 14 | Yes |
| 7 |  | >5 | 24 | 25 | 12 | Yes |
| 8 |  | 2 | 21 | 22 | 11 | No<sup>a</sup> |
| 9 |  | >5 | 26 | 22 | 8 | No<sup>b</sup> |
| 10 | HS11 | >5 | 47* | 47* | 0* | Not sequenced |
| 11 |  | 2 | 0* | 25 | 25* | Yes |
| 12 |  | 2 | 31* | 25 | 7* | Yes |
| 13 |  | 3 | 32* | 20 | 6* | No<sup>b</sup> |
| 14 | HS4 | 1 | 26 | 0 | 0 | N/A |
| 15 |  | 2 | 40* | 0 | 0 | N/A |
| 16 |  | 1 | 18 | 0 | 0 | N/A |

\* Indicates deviation from expected segregation of single locus or mono-allelic gene targeting, N/A: Not applicable (GT negative control)
<sup>a</sup> Deletion of partial ZMUBI intron of donor template
<sup>b</sup> Duplication of HR2 homology sequence at 3' end of the donor template

## Discussion

Gene targeting relies on endogenous homologous recombination machinery, which is used to repair genomic DNA damage due to DSBs in somatic cells or during meiosis for crossovers between homologues^35^. DSBs activate the cell’s DNA repair machinery making the break site accessible to a donor template for potential GT^36–38^. Due to flexibility and ease of the design, CRISPR-Cas9 has become an important tool for generating targeted DSBs for genome engineering applications^1,39–41^. In somatic cells, NHEJ, which mainly involves ligation of unrelated sequences or sequences with micro-homologies, is the primary DNA repair pathway^7^. While targeted mutagenesis using CRISPR-Cas9 is now routine in plants that are amenable for transformation, HR-mediated targeted insertion or allele replacement remains a challenge^42,43^. The low frequency of recovering GT events is further hindered by the general dependence on a selectable marker to enrich for GT within the population of recovered plants. Development of a selectable marker-free GT system is highly desirable for downstream application and product deployment.

In this report we demonstrate high frequency GT in maize, by relying on Cas9-mediated release of a marker-free donor template leaving the activated by the DNA repair selectable marker behind in the original randomly-integrated T-DNA locus. Given that delivery of DSB reagents along with donor template and accessibility at the DSB site are key bottlenecks, we applied our High-IGHR method that utilizes randomly pre-integrated donor template and CRISPR reagents. Taking advantage of the plant’s inherent cell division processes, the cells containing donor template and CRISPR reagents are enriched and later utilized to release donor template for intra-genomic GT. Intra-genomic homologous recombination has previously been demonstrated in whole plants ^22,28–30^. Recent improvements in GT frequency have used either different CRISPR systems^44^, an egg cell-specific promoter driving nuclease expression^45^, or sequential transformation^46^. These recent developments have been demonstrated mainly in *Arabidopsis*. Moreover, all these reports rely on low frequency GT coupled with extensive selection and/or screening over multiple generations, making these approaches impractical for routine application in long life cycle crop plants.

To obtain stable, non-chimeric, IGHR-mediated GT in a single generation, we hypothesized that delayed induction of Cas9 following plant transformation should be adequate to provide CRISPR reagent- and donor template-enriched cells. In addition, Cas9 induction and GT in undifferentiated cells would ensure regeneration of non-chimeric, GT plants from single cells. Since our goal was to develop a GT system without a selectable marker in the donor template, regeneration that enriched for rare targeted cells was the next major hurdle. Although our donor template contained the *NptII* expression cassette, no corresponding selection (e.g. with G418) was applied throughout the experiment thus alleviating the need of selectable marker in the donor template. Instead, we designed vectors where the donor repair template flanked by Cas9 target sites was positioned between the *Sb-ALS* promoter and *Hra* coding sequence, thus precluding expression of the selectable marker gene while in this configuration. Following Cas9 induction, release of the donor repair template and subsequent NHEJ-mediated repair of the donor locus re-constituted the functional selectable marker expression cassette. While this approach does not directly permit selection of cells with GT, the selective advantage for the Cas9-expressing cells with donor release provided preferential enrichment for GT events.

Accordingly, the heat shock-inducible Cas9 and selectable marker activation system demonstrated preferential regeneration of repair template-released T0 plants, as 86% of those plants were PCR-positive for donor template release. The mutation analysis of target site revealed that up to 99% of T0 plants contained mutations in at least one allele, while >75% of the plants were bi-allelic mutants. Diagnostic junction PCR analysis of these plants revealed up to 4.7% GT, which is similar to the GT frequency previously reported using a donor repair template that contained a selectable marker delivered via particle bombardment^44^. GT without a selectable marker in the repair template has been previously reported in maize events produced via particle bombardment^19^; however, the observed frequencies (0.5-1%) were lower than those observed in this report. We attribute this roughly 5-fold enhancement in GT frequencies to our new High-IGHR method.

One intriguing observation in this study was the differences in the number of plants obtained when comparing two heat shock treatments applied to induce Cas9 expression. Approximately 2.5-fold more plants were regenerated when Cas9 was induced 4 days (HS4) after *Agrobacterium* infection compared to induction after 11 days (HS11) (Table 1). While there was 7-day resting period between Cas9 induction and selection for HS4 treatment, the explants from HS11 group were transferred to selection media without any resting on non-selective media. We believe that elimination of the resting period between Cas9 induction (with resultant *Hra* activation) and transfer to selective media could have contributed to the lower number of plants regenerated from the treatment HS11 (i.e., multicellular HRA-expressing cell clusters (HS4 treatment) were more resistant to the selective pressure than single cells (HS11 treatment). Further, higher frequency GT (4.7%) was observed in the HS11 group with later Cas9 induction in comparison to HS4 group (2.7%). Based on these observations, we speculate that GT frequency could be increased by extending resting period post-Cas9 induction in the HS11 treatment. Compared to HS4 treatment, explants in HS11 group were enriched for cells containing Cas9 reagents and donor template (potentially 500-1000-fold)^45^ due to the additional 7 days of growth prior to Cas9 induction. Refinement of the time interval of Cas9 induction and donor template release followed by stringent selection regime for plant regeneration could further improve GT. While the inducible intra-genomic approach applied here enriches for cells containing the GT reagents, the low number of donor template copies per cell remains a key limitation, which could be addressed by using viral replicon system^46^ to further enhance GT.

After having achieved high frequency GT in T0 plants, the next goal was to validate that GT was non-chimeric and stably inherited. Analysis of T1 progeny of 13 GT-positive T0 plants revealed Mendelian mono-allelic inheritance and segregation of GT-modified target locus for 11 plants, while two plants showed bi-allelic GT. These observations confirm our hypothesis that controlled induction of Cas9 during a later undifferentiated stage of tissue culture generates non-chimeric, heritable GT events. Compared to previous IGHR reports ^22,28–30^, this is a major advancement both for improved GT frequency and generation of non-chimeric GT plants in a single generation.

Sequencing of the junction PCR products revealed approximately 600 bp deletion from the donor sequence in T1 plants from two T0 plants, and a duplication of truncated HR2 3’-end sequence in T1 plants from three T0 plants. The highly repetitive nature of the ZMUbi promoter intron sequence^47^ could have contributed to the deletion. Both events containing the HR2 3’end duplication derived from the same immature embryo and potentially could be clonal. Additionally, the HR2 3’-end fragment observed in three T0 plants could be attributed to the one-sided invasion model of recombination^48^, which suggests that homology to only one end of the DSB is sufficient for HR.

In summary, we have developed a high frequency, selection-free GT system in a major crop plant. Although to test this method we used our transformation-competent process with the morphogenic genes *Wus2* and *Bbm*, the approach should be amenable and perhaps even more useful in crops or genotypes that are recalcitrant to transformation or transform at impractically low frequencies. A handful of transgenic cultures could be enriched in undifferentiated tissue culture stage for IGHR to obtain selection-free GT in a single generation. In addition, the selectable marker activation system described here would be an attractive design for improving frequency of recovering GT, targeted mutagenesis or deletions in plant genome.

## Methods

### Plant material

Maize (*Zea mays* L.) inbred line PH1V69 was obtained from internal Corteva Agriscience sources.

### Plasmids and reagents used for plant transformation

Cas9 and guide RNA vector construction was previously described in Svitashev et al. (2015)^44^. The morphogenic transcription factors *Babyboom* (*Bbm*), and maize *Wuschel2* (*Wus2*), which in the process of stimulating somatic embryogenesis also promote cell division, have also been described previously^49^. Supplementary Table 1 provides the details of selectable markers and other elements used in different constructs described in Figure 2.

### Maize transformation

Corteva Agriscience inbred line PH1V69 was grown in greenhouse growth conditions. Ear harvest, immature embryos isolation, *Agrobacterium*-mediated transformation, and plant regeneration were based on those described by Jones et al. (2019)^34^.

### Intra-genomic GT using inducible Cas9

For T0 GT plant production, three treatments were performed after *Agrobacterium* infection (refer to Figure 3). For the first treatment (HS4) the heat shock (45 °C for 3 hours with 70 % RH) was applied 4 days after *Agrobacterium* infection for 2 consecutive days (heat treatment on day 4 and again on day 5) followed by 7 days on Resting Medium (RM) with no selection. Twelve days after *Agrobacterium* co-cultivation, the embryos were transferred to RM with 0.1 mg/L Ethametsulfuron (Sigma-Aldrich^®^; St. Louis, MO) for 2 weeks followed by 2 weeks on Maturation Medium (MM) with 0.05 mg/L Imazapyr. The rooting step was performed on Rooting Medium (RtM) containing 0.05 mg/L Imazapyr for 2-3 weeks. The second treatment (HS11) is identical to HS4 with the main difference being the timing of the first heat shock that is applied 11 days after *Agrobacterium* infection (again on two consecutive days 11 and 12), and then transferring immediately to RM with 0.1 mg/l Ethametsulfuron. Subsequent culture steps for the HS11-treated material were identical to the HS4 method. For the third treatment (Control), no heat shock was applied to the immature embryos. These embryos followed the same tissue culture and regeneration steps as HS4 and HS11 treatments. Once roots have formed, plantlets were transferred to the greenhouse. Pollen from the T0 plants was used for cross pollination of wild type PH1V69 ears to obtain backcross 1 (BC1) progeny.

### Molecular analysis

DNA extraction from maize tissue and copy number qPCR was completed for T-DNA left and right borders along with *NptII* selection element as described in Anand et al. (2019)^50^. (LB F-CATGAAGCGCTCACGGTTACTAT, LB R-TCGTACGCTACTGCCACCAA, LB Probe-6FAM-ACGGTTAGCTTCACGACT) (RB F-TGATTCCGATGACTTCGTAGGTT, RB R-GCTAATCGTAAGTGACGCTTGGA, RB Probe-6FAM-CTAGCTCAAGCCGCTC) (NPTII MO1 F-GACCTTTTGTCCAGCCATCTG, NPTII MO1 R-GGCGTCTGCCATGATGCT, NPTII MO1 Probe-6FAM-CACCTGCCGAAAAG). Mutation frequencies at genomic target sites were analyzed by qPCR using primer pair (F-GTAACCTACAAGGCCACGGACTG, R-TAGGACGGCCACAATCAAAGTT) and probes (6FAM-TGCTCCTCATCCTCGAT, VIC-TCTAGCTTTGCATCATGTC). To verify donor template release from the genomic integration site, PCR amplification was performed using a primer pair annealing to the *Sb-ALS* promoter (CATCAACCGGCTCTCCCTTTC) and *Hra* coding region (GACGTCGAGGACCAGGTAGTTGT). The PCR reaction with the described primers could amplify either 5374 bp fragment if donor template is not released or a 670 bp fragment indicating dropout of the donor template. The PCR conditions were set up to amplify only the short 670 bp product using REDExtract-N-Amp PCR ReadyMix (Sigma). PCR conditions: DNA denaturation at 94°C for 5 min, 94°C for 30 sec, annealing at 62°C for 30 sec, extension at 72°C for 1 min, 35 cycles. For diagnostic GT analysis, the presence HR1 and HR2 junctions (Figure 3B) were analyzed by qPCR using HR-spanning primers (F1 HR1-GCGTGCGTGCTTACATGATG genomic, R1-AGACATGCAATGCTCATTATCTCTAG insert, F2 HR2 – AAACAGATAAACGGGACTCTCCAG insert, R2-GTGCGACATTAAACAGTGTTAGTTGTAGCC genomic), followed by nested primers (nested HR1 F1-TGCAGTGCAGCGTGACC, nested R1-AGACATGCAATGCTCATTATCTCTAG, probe 6FAM-AGAGGGGCACGACC) (HR2 F1-GCGTGCGTGCTTACATGATG genomic, R1-GCTTGAAGCCTACCAAAGCAA and probe 6FAM-AATGATCGACAAACGCT). To further verify the integrity of the GT events, long overlapping PCR (Figure 3C and D) was performed with primer pairs (HR1 F1-GCGTGCGTGCTTACATGATG genomic, R1-AAAGTTCAGAATAGATTTGGTTTGGA insert) and (HR2 F2-CATTTTATTAGTACATCCATTTAGGGTTTAG insert, R2-GTGCGACATTAAACAGTGTTAGTTGTAGCC genomic) using NEB LongAmp^®^ Hot Start Taq 2X Master Mix (Catalog# M0533S) according the manufacture’s recommendations. The long HR spanning PCR products were then sequenced using standard Sanger sequencing method.

## Authors’ contributions

S.K., P.B., S.S., A.A, and W.G.K. designed the experiment. E.W. and P.B. performed the maize transformation. B.L. designed and performed DNA analysis. S.K. led the project and wrote the manuscript. All the authors reviewed the manuscript.

## Acknowledgements

We thank Larisa Ryan, Zhi Li, Julia Chow-Yiu, Ying Yan, Spencer Jones, Anne Ulm, Grace St Clair, and Lanie Feigenbutz for maize transformation, and Craig Hastings and Lijuan Wang for vector construction. The authors also thank Scott Betts, Maria Fedorova, Sendil Devadas, and Tracey Fisher for critical review of the manuscript. We acknowledge Clara Alarcon and Doane Chilcoat for providing resources and support for this work.

## Data availability

The data supporting the findings of this study are available within the paper and its supplementary information files.

