## Supplemental Table 1 for "Efficient Gene Targeting in Maize using Inducible CRISPR-Cas9 and Marker-Free Donor Template"

**Supplementary Table 1**. Description of the elements used in the T-DNA vector shown in Figure 2.

| Features | References |
| --- | --- |
| *HSP26 & HSP17.7* | ^1^ |
| *Cas9, U6 promoter, and guide RNA* | ^2^ |
| *SB-ALS Pro* | SB-ALS promoter and 5’UTR, DOE-JGI Sbi v3.1, SBChr04, bases 49239164-49240031. DOE-JGI Sbi v3.1 corresponds to Sorghum bicolor BTx623 assembly v3.0.1 and gene annotation v3.1 available from phytozome (http://phytozome.jgi.doe.gov/). Chromosome 4 of Sbi v3.1 is registered as NCBI accessions NC_012873.2 and CM000763.3 |
| *HR1* | ^3^ |
| *HR2* | ^3^ |
| *Cas9 cut site* | ^3^ |
| *ZMUbi-nptII* | ^4^ |
| *HRA* | ^5^ |
| *Wus2 & Bbm* | ^6^ |

1	Anand, A. *et al.* Methods and composition for rapid plant transformation. US patent 2170121722 A1 (2017).

2	Svitashev, S. *et al.* Targeted Mutagenesis, Precise Gene Editing, and Site-Specific Gene Insertion in Maize Using Cas9 and Guide RNA. *Plant Physiol* **169**, 931-945, doi:10.1104/pp.15.00793 (2015).

3	Cigan Andrew, M. *et al.* Generation Of Site-specific-integration Sites For Complex Trait Loci In Corn And Soybean, And Methods Of Use. WO patent WO 2016/040030 A1 (2016).

4	Anand, A. *et al.* High efficiency Agrobacterium-mediated site-specific gene integration in maize utilizing the FLP-FRT recombination system. *Plant Biotechnol J*, doi:10.1111/pbi.13089 (2019).

5	Green, J. M., Hale, T., Pagano, M. A., Andreassi, J. L. & Gutteridge, S. A. Response of 98140 Corn with gat4621 and hra Transgenes to Glyphosate and ALS-Inhibiting Herbicides. *Weed Sci* **57**, 142-148, doi:Doi 10.1614/Ws-08-152.1 (2009).

6	Lowe, K. *et al.* Morphogenic Regulators Baby boom and Wuschel Improve Monocot Transformation. *Plant Cell* **28**, 1998-2015, doi:10.1105/tpc.16.00124 (2016).
